# Regulation by the Pitcher Plant Sarracenia purpurea of the Structure of its Inquiline Food Web

**DOI:** 10.1101/2020.05.15.098004

**Authors:** Aaron M. Ellison, Nicholas J. Gotelli, Leszek A. Błędzki, Jessica L. Butler

## Abstract

Phytotelmata, the water-filled habitats in pitcher plants, bromeliad tanks, and tree-holes, host multitrophic food webs that are model experimental systems for studying food-web structure and dynamics. However, the plant usually is considered simply as an inert container, not as an interacting part of the food web. We used a manipulative field experiment with a response-surface design to determine effects of nutrient enrichment (multiple levels of NH_4_NO_3_, PO_4_, and captured prey), top predator (removed or present), and the plant itself (with or without plastic tubes inserted into the pitchers to isolate the food web from the plant) on the macrobial food web within the modified leaves (“pitchers”) of the carnivorous pitcher plant *Sarracenia purpurea*. Connection to the plant, addition of NH_4_NO_3_ and removal of the top predator significantly increased the food web’s saturation, defined as its trophic depth and number of interactions. No effects on food-web saturation resulted from addition of PO_4_ or supplemental prey. Plants such as *S. purpurea* that create phytotelmata are more than inert containers and their inhabitants are more than commensal inquilines. Rather, both the plant and the inquilines are partners in a complex network of interactions.

## Introduction

Understanding controls on food web structure has been an organizing focus for community ecology for well over a century (*e.g.*, Pimm, 1982; Polis, 1994; Carpenter *et al.*, 2001; McCann, 2012; Wokovich *et al.* 2014; Layman *et al.*, 2015; Zou *et al.*, 2016). As primary producers, plants are the bottom-up controllers of “green” food webs, whereas as “detritus” the decomposing leaf litter of plants is the resource base for “brown” food webs (Cebrian and Lartigue, 2004). The bottom-up roles of plants as producers of fresh biomass and necromass in structuring, respectively green and brown food webs, have been studied extensively, but other roles for plants in structuring food webs through habitat creation and indirect interactions is much less appreciated. For example, in tree holes and modified leaves, plants create habitats for a range of aquatic food webs. The complex, mostly detritus-based food webs in these “phytotelmata”—water-filled habitats in, for example, tree holes, bromeliads, and pitcher plants (Maguire, 1971)—have been used as model experimental systems for studying the dynamics of aquatic food webs (*e.g.*, Kitching, 2000, 2001; Kneitel and Miller, 2002; Miller *et al.*, 2002; Ellison *et al.*, 2003; Srivastava *et al.*, 2004). However, the phytotelmata itself normally is regarded simply as a container for its “inquilines”: animals that live commensally “within the nest or abode of another” (OED, 2019). It has been recognized only rarely that the plant itself can mediate trophic interactions of inquiline food webs.

Research with carnivorous pitcher plants has provided two notable examples of the importance of phytotelmata on inquiline food webs. Bradshaw and Creelman (1984) experimentally showed that the carnivorous pitcher plant *Sarracenia purpurea* L. (Sarraceniaceae) and two of its obligate inquilines, larvae of the pitcher-plant mosquito *Wyeomyia smithii* (Coq.) and the pitcher-plant midge *Metriocnemus knabi* Coq., are mutualists. These two inquilines accelerate the breakdown and decomposition of arthropod prey captured by *S. purpurea* (Heard, 1994). Ammonium (NH_4_) and carbon dioxide (CO_2_) released as bacteria mineralize nutrients in the prey (Butler *et al.*, 2008) and NO_3_, NH_4_, and PO_4_ regenerated by rotifers (Błędzki and Ellison, 1998, 2002; Błędzki *et al.*, 2018) are absorbed by the plant and used for photosynthesis, subsequent growth, and reproduction (Bradshaw and Creelman, 1984; Joel and Gepstein, 1985; Butler and Ellison, 2007). The plant, in turn, oxygenates the pitcher fluid (Bradshaw and Creelman, 1984; Joel and Gepstein, 1985; Sirota *et al.*, 2013), which maintains homeostatic conditions for the inquiline food web and prevents a switch to anaerobic conditions within the phytotelmata, which can nonetheless occur if prey are superabundant (Krieger and Kourtev, 2012; Sirota *et al.*, 2013; Northrop *et al*., 2017).

Similarly, one of the Asian pitcher plants, *Nepenthes bicalcarata* Hook.f. (Nepenthaceae) hosts the ant *Colobopsis schmitzi* (Stärcke) (Formicidae) in the plant’s hollow tendrils (Clarke and Kitching, 1995). The ants move out of the tendrils to forage within the pitchers on large prey that the plant captures. The ants also clean the pitcher’s peristome, reducing fungal colonization and increasing prey-capture efficiency (Thornham *et al.*, 2011), which in turn leads to greater nutrient acquisition, photosynthesis, and growth of this perennial plant (Bazile *et al.*, 2012).

The lack of attention to linkages between food-web structure or dynamics and the habitat in which the food web itself assembles is not restricted to phytotelmata. Although spatiotemporal dynamics of food webs have been studied, documented, and modeled (*e.g.*, Poisot *et al.*, 2015; Pilosof *et al.*, 2017; Piovia-Scott et al. 2017; Hutchinson et al., 2019; Schiaffino *et al.*, 2019), how complete food webs assemble in a habitat that itself is changing through time and interacts with the food web itself remains unexplored. We are unsure whether this lack of attention results from the rarity of such situations, our failure to perceive them, or the difficulty in studying them. However, living phytotelmata such as pitcher plants provide opportunities to study experimentally the dynamics of a food web that assembles in a habitat that itself is a living, growing organism. The experiments we describe here were designed to test whether pitcher plants are more than simply “containers” for “inquilines” and how the interactions between plants, the habitat it creates, and the food web that assembles within it change through time.

Here, we describe how nutrient enrichment, top predators, and the living habitat of the plant itself affect the structure of the four trophic-level macrobial food web that inhabits the modified pitcher-shaped leaves (Arber, 1941) of *S. purpurea*. In a set of controlled field experiments, we first examined the impact of nutrient deposition on the food web that assembles in *S. purpurea* pitchers. Second, we severed the connection between the plant and its food web by inserting closed plastic tubes (or appropriate controls) in between pitchers and their food webs to determine if effects of experimental nutrient enrichment through additions of prey and inorganic nutrients resulted from direct effects on the food web or indirect effects mediated by the plant. We recognize that a focus on the macrobes will overlook many important processes and interactions of the microbes present in *S. purpurea* food webs. This is a general issue with studies of food webs (e.g., Pringle and Hutchinson, 2020) that will be able to be addressed as the microbial assemblages of *S. purpurea* continue to be better characterized (e.g., Peterson *et al.* 2008, Miller and terHorst 2012, Paise *et al.* 2014, Canter *et al.* 2018, Boynton *et al.* 2019, Zhang *et al.* 2020a, 2020b).

## Materials and Methods

### Study Species

*Sarracenia purpurea* (the northern, or purple, pitcher plant) is a long-lived (> 50 yr), rosette-forming, carnivorous perennial plant common to bogs throughout Canada and the eastern United States. Pitcher plants flower early in the growing season (May in Massachusetts) before any new leaves are produced; flower buds for the following season are set in August – September (Shreve 1906). Pitcher plants have weakly developed root systems; nutrients are obtained principally from insect prey attracted to the brightly colored pitchers and extrafloral nectaries (Bennett and Ellison, 2009). Unlike the other 10 species of *Sarracenia* (Mellichamp and Case, 2009)*, S. purpurea* pitchers fill with rainwater and these semi-permanent pools (phytotelmata) are colonized by a suite of specialized invertebrates and more generalist protists and bacteria (Addicott, 1974; Cochran-Stafira and von Ende, 1998; Błędzki and Ellison, 2003; Ellison *et al.*, 2003, Peterson *et al.* 2008, Miller and terHorst 2012, Paise *et al.* 2014, Canter *et al.* 2018, Boynton *et al.* 2019, Zhang *et al.* 2020a, 2020b).

Arthropod prey captured by the plant is the basal resource for this detritus-based food web, which shreds the prey and mineralizes the available nutrients (Butler and Ellison, 2007; Butler *et al*., 2008). Prey capture initially increases as pitchers open, expand, and harden; reaches a peak when the leaves are two–three weeks old; and then declines during to near zero by the end of the growing season (Fish and Hall, 1978; Ellison *et al.*, 2003). Drowned carcasses of captured prey (primarily ants and flies; Ellison and Gotelli, 2009) are shredded by larvae of the pitcher-plant midge *Metriocnemus knabi* and larvae of the sarcophagid fly *Fletcherimyia fletcheri* (Aldrich); the shredded prey are broken down further by bacteria (Cochran-Stafira and von Ende, 1998; Heard, 1994; Baiser *et al.*, 2011). The bacteria, organic matter, protozoa, and algae in turn are consumed by the bdelloid rotifer *Habrotrocha rosa* Donner and a variety of protozoa, which, along with the pitcher-plant mite *Sarraceniopus gibsoni* (Nesbitt), also consume detritus directly (Wallace *et al.*, 2006). Protozoa also consume bacteria (Cochran-Stafira and von Ende, 1998). These primary consumers are preyed on by larvae of the pitcher-plant mosquito *W. smithii* (all these macrobes will be referred to below using their generic epithets). Late instars of *Fletcherimyia* also prey on rotifers and 1st and 2nd instars of *Wyeomyia* (Błędzki and Ellison, 1998; Butler *et al.*, 2008; Baiser *et al.*, 2013). Not only does *Fletcherimyia* complete its larval development within pitchers, but the adult flies are pollinators of *S. purpurea* (Ne’eman *et al.*, 2006).

### Study Site

The experiments reported here were conducted at Swift River Bog, a 1.9-ha glacial kettle bog in Belchertown, Massachusetts, USA (42.27 °N, −72.34 °W, 124 m a.s.l.; Kearsley, 1999). This bog has a lagg (moat) around it that never exceeds 2 m in width, but there is no open water within the bog itself. Common *Sphagnum* species present at Swift River Bog were *Sp. angustifolium* (C.E.O. Jensen ex Russow) C.E.O. Jensen*, Sp. magellanicum* Brid., and *Sp. rubellum* Wilson (Ellison and Gotelli, 2016). The dominant vascular plants growing on the bog mat were shrubs in the Ericaceae: *Chamaedaphne calyculata* (L.) Moench (leatherleaf)*, Kalmia polifolia* Wangenheim (bog laurel), *K. angustifolia* L. (sheep laurel), *Rhododendron viscosum* (L.) Torrey (swamp rhododendron) and *Vaccinium macrocarpon* Aiton (cranberry); scattered *Picea mariana* (Miller) Britton (Pinaceae; black spruce) grow across the bog. The site has a large population (1000s of plants) of *S. purpurea* (Ne’eman *et al.*, 2006; Ellison and Gotelli, 2016). Total nitrogen deposition (NO_3_ + NH_4_) at the National Atmospheric Deposition Program (NADP) Quabbin Reservoir monitoring station in the same watershed (MA08: 42.3925 °N, −72.3444 °W) ranged from 1.25–1.60 mg/L during the years of this study (2001 – 2004), and has averaged 1.19 (range: 0.58–2.12) mg/L since measurements began in 1982 (National Atmospheric Deposition Program, 2020). Phosphorus deposition is not monitored by NADP or other agencies in the US, but P deposition may have significant effects in P-limited ecosystems or alter stoichiometric ratios in otherwise N-limited ones (Mahowald *et al.*, 2008; Tipping *et al.*, 2014). Although *S. purpurea* is stoichiometrically N- or N+P-limited (Ellison, 2006), field experiments have suggested that *S. purpurea* preferentially takes up P from supplemental prey (Wakefield *et al.*, 2005).

### Food-Web Saturation: Definition of the Response Variable

In all the experiments described below, *food-web saturation* was our response variable. We defined food-web saturation as follows: Each macrobial taxon (rotifer, mite, midge, mosquito, sarcophagid fly) in the food web was assigned a binary value representing its presence (Table 1; Fig. 1); we assumed all food webs had protozoa and bacteria. The saturation of the food web in a pitcher was defined as the sum of the equivalent decimal values of each macrobial taxon present. There are 32 possible food webs that can be assembled from these five taxa; the decimal value for each food web ranges from 0 (no macrobial taxon present) to 31 (all macrobial taxa present), with increasing numbers indicating more “saturated” food webs. Food webs with higher saturation values had both more trophic levels present and more trophic links present (Fig. 1), but the food-web saturation index does not reflect any particular trophic relationship. Further, the index can be easily extended if any other taxa (e.g., microbes) were to be included.

**Table 1.** Values assigned to each inquiline taxon for calculation of a food-web saturation index. Occurrence of taxa in a given food web is given by the sum of the binary values of all taxa present. The saturation index is the sum of the decimal values of all taxa present. See Fig. 1 for an illustrative example of these calculations

| **Taxon** | **Binary**<br>**value** | **Decimal**<br>**value** |
| --- | --- | --- |
| *Metriocnemus* | 1 | 1 |
| *Habrotrocha* | 10 | 2 |
| *Sarraceniopus* | 100 | 4 |
| *Wyeomyia* | 1000 | 8 |
| *Fletcherimyia* | 10000 | 16 |

**Fig. 1.**
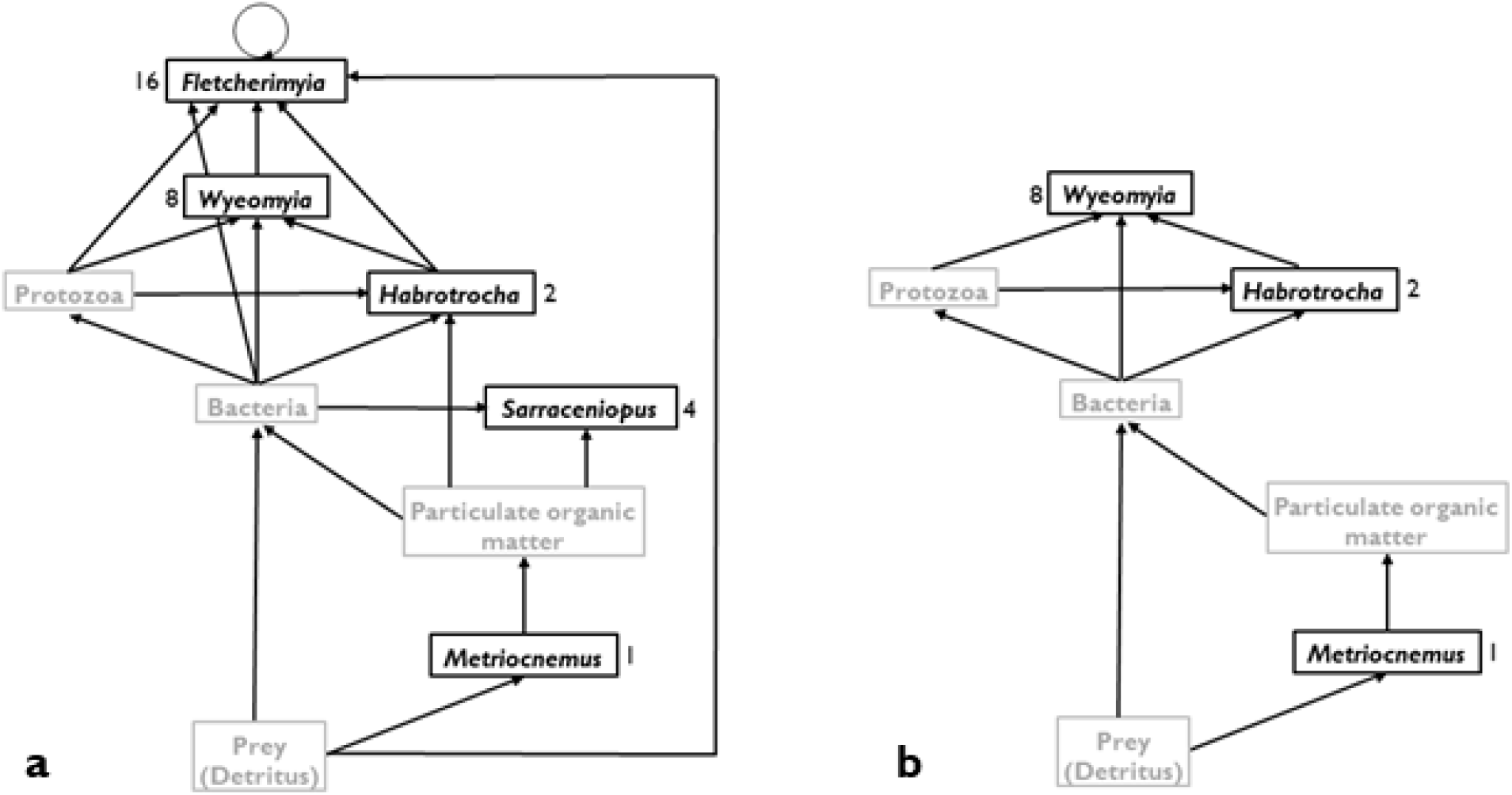
Complete, fully saturated inquiline food web of *Sarracenia purpurea* (**a**; modified from Baiser *et al.*, 2013) and an example of incomplete, unsaturated one (**b**). Black type indicates the “macrobes” we studied here; grey type the microbes and detritus that we did not quantify. Values adjacent to each of the macrobial taxa are those used to compute food-web saturation (see Table 1). Prey, bacteria, protozoa, and particulate organic matter were assumed to occur in all sampled food webs, and each of the possible combinations of macrobes gives a unique saturation value. In these two examples, the saturation value of the complete web (a) = 31, whereas that of the food web lacking *Fletcherimyia* and *Sarraceniopus* (b) = 11

### Experiment 1 – Impacts of N and P on Food-Web Saturation

The first experiment investigated the effects of inorganic N and P (as would be accumulated in the phytotelmata by atmospheric deposition) on food-web saturation in situ. We used a factorial response-surface design (Cottingham *et al*., 2005) in a “pulse” experiment (sensu Bender *et al*., 1984) in which 5 levels each of NH_4_NO_3_ (4.687, 9.375, 18.75, 37.5, and 75.0 mM) and PO_4_ (0.625, 1.25, 2.5, 5.0, and 10.0 mM) and controls (distilled, deionized water: dH_2_O) were added biweekly to each pitcher of 130 non-flowering plants that were randomly assigned to one of five groups of 26 plants. The levels of N used bracketed the observed deposition of N measured by NADP in the years before the study, whereas the levels of P were chosen to span the N:P ratios previously used by Ellison and Gotelli (2002) in other nutrient addition experiments with *S. purpurea* without any evidence of nutrient toxicity (*sensu* Camargo *et al*. 2005).

Plants were located at least 10 m (linear distance) from the lagg surrounding the bog, separated from one another by at least 50 cm, and atop the *Sphagnum* mat but not under shrubs or trees. Each plant within a group was randomly assigned a nutrient treatment (one of the 25 possible N × P combinations or the dH_2_O control). On 4 June 2001, 20 ml of nutrient solution was added once to each open pitcher on each plant. Plants were monitored every three d for new pitcher production, and nutrient solutions were added (again, only once) to each new pitcher 2-3 days after it had opened and fully hardened. Every 3 wk (25 June, 16 July, 6 August, 27 August, and 17 September), a complete set of 26 plants was harvested. At each harvest, pitcher fluid was pipetted into sterile tubes (one tube/leaf), the leaves further rinsed to ensure that all rotifers and mites were collected, and the presence or absence of each member of the food web was determined.

We ran a parallel “press” experiment (sensu Bender *et al.*, 1984) in which we haphazardly located 26 non-flowering plants and randomly assigned each of them to one of the same 26 nutrient treatments as in the pulse experiment. 2-3 d after each pitcher had opened and hardened, 20 ml of nutrient solution was added to it, and the 20-ml level was marked on the side of the pitcher with indelible ink. Every two weeks, the food web was pipetted out of each pitcher and into a petri dish; the volume of the flued was recorded. Presence of each macrobe was determined in the field, and all were returned to the pitcher. Rotifers and protozoa were counted in the lab in 1-ml subsamples taken at the same time; total counts of rotifers and protozoa were extrapolated to the full volume by simple multiplication. After field counts of invertebrates, the pitcher fluid was poured back into the pitcher and additional nutrient solution was added to maintain the fluid level at 20 ml. This experiment ran from 4 June–17 September 2001.

### Experiment 2 – Interactions Between the Plant, Nutrients, and Predation on Food-Web Structure

The second experiment investigated how changes in food-web saturation resulted from interactions between bottom-up effects of nutrient addition, top-down effects of predation, and effects of the plant itself. We used a four-level factorial/response-surface experiment with 212 non-flowering plants located at least 10 m (linear distance) from the lagg surrounding the bog, separated from one another by at least 50 cm, and atop the *Sphagnum* mat but not under shrubs or trees. As in Experiment 1, plants were assigned randomly to unique combinations of treatments that included physically isolating the inquiline food web within the phytotelmata.

The first treatment (factor) assessed the influence of the plant itself on the food web and had three levels. Pitcher interiors were: [1] fitted with intact polyethylene tubes (2.2-cm diameter) (inquilines and their fluid isolated from effects of pitchers; n = 64 plants, 1 pitcher/plant); [2] fitted with perforated (ten 6.35-mm diameter drilled holes) polyethylene tubes (polyethylene tube control; n = 64 plants); or [3] left without polyethylene tubes (unmanipulated control; n = 64 plants).

The second treatment (ordinal factor used in the response surface) examined the effects of N deposition. One of eight different concentrations of inorganic nitrogen (NH_4_NO_3_: 0, 1.56, 3.1, 6.25, 12.5, 25, 50, or 100 mg N/L) was added to a randomly selected plant; there were n = 8 replicates of each N concentration in each set of 64 tube/tube-control/no tube treatments. Because we assumed a constant uptake of NH_4_NO_3_ by pitcher tissue in control and tube-control pitchers, and volatilization and uptake by bacteria in all pitchers, we replaced fluid and nutrients with the appropriate NH_4_NO_3_ solution every two weeks.

The third treatment (factor) examined effects of prey availability (organic nutrients). Supplemental prey (10 lab-raised fruit flies/wk) were added to one-half of each inorganic N × tube combination.

Finally, because of the importance of *Fletcherimyia* at multiple points in this food web (Błędzki and Ellison, 1998; Butler *et al.*, 2008; Baiser *et al.*, 2013), the fourth treatment (factor) assessed the influence of this top predator on the development of the pitcher-plant food web in the context of bottom-up effects of prey and inorganic N addition and habitat effects of the pitcher itself. Second instar *Fletcherimyia* larvae were either removed from pitchers (one-half of each N × tube × fruit-fly treatment combination) or added to pitchers (one-half of each N × tube × fruit-fly treatment combination).

In total, this factorial/response-surface design resulted in *N* = 2 plants in each tube × N × fruit-fly × *Fletcherimyia* treatment combination. We also haphazardly chose 20 additional plants growing nearby in the bog as unmanipulated controls. One treated plant from each treatment combination (96 pitchers) and half the unmanipulated controls were harvested after 4 wk, and the remainder after 8 wk. Food-web saturation in this experiment was computed based on all inquilines present including *Fletcherimyia*, except in the *Fletcherimyia* addition treatments (to avoid simply documenting the addition in our calculation).

### Statistical Analysis and Data Availability

Data were analyzed using standard linear regression, analysis of variance, or analysis of covariance using functions for generalized linear models (glm) in the base and stats packages of the R statistical software system, version 3.6.3 (R Core Team 2020).

Experiment 1 (pulse and press experiments) were analyzed using a linear regression model (i.e., linear in the regression parameters) with which we fit a polynomial surface of food-web saturation as function of N and P concentrations and their interaction. Experiment 2 (four treatment factorial/response-surface design) was analyzed using a glm with main effects of the tubes (3 levels), supplemental prey (2 levels), *Fletcherimyia* addition (2 levels), inorganic N addition (8 ordered levels treated as a continuous variable in the model), and all of their two- or three-way interactions. Initial analyses revealed no significant interaction effects among the three discrete factors (P > 0.20 all cases), so we report only the results of the main effects and the interactions with the inorganic N additions. Throughout, F statistics and associated P values are based on Type III sums of squares.

All data from this experiment are available from the Harvard Forest Data Archive, dataset HF193 (https://harvardforest.fas.harvard.edu/exist/apps/datasets/showData.html?id=hf193) and the Environmental Data Initiative (https://dx.doi.org/10.6073/pasta/3bd2c920cf431442d34227f3f5fc7c2a).

## Results

### Experiment 1

In the pulse experiment, which established the baseline response of the inquiline food web to bottom-up control by inorganic nutrient additions, food-web saturation peaked at intermediate levels of N added (F_1,227_ = 10.7. P < 0.01), was significantly higher in pitchers produced earlier in the season (F_1,227_ = 5.0, P = 0.03), and in leaves harvested in mid-summer (F_1,227_ = 4.4, P = 0.04) (Fig. 2). Although the amount of P added did not alter food web saturation (F_1,227_ = 0.7, P = 0.42), there was a significant N × P interaction (F_1,227_ = 4.4, P = 0.04).

**Fig. 2.**
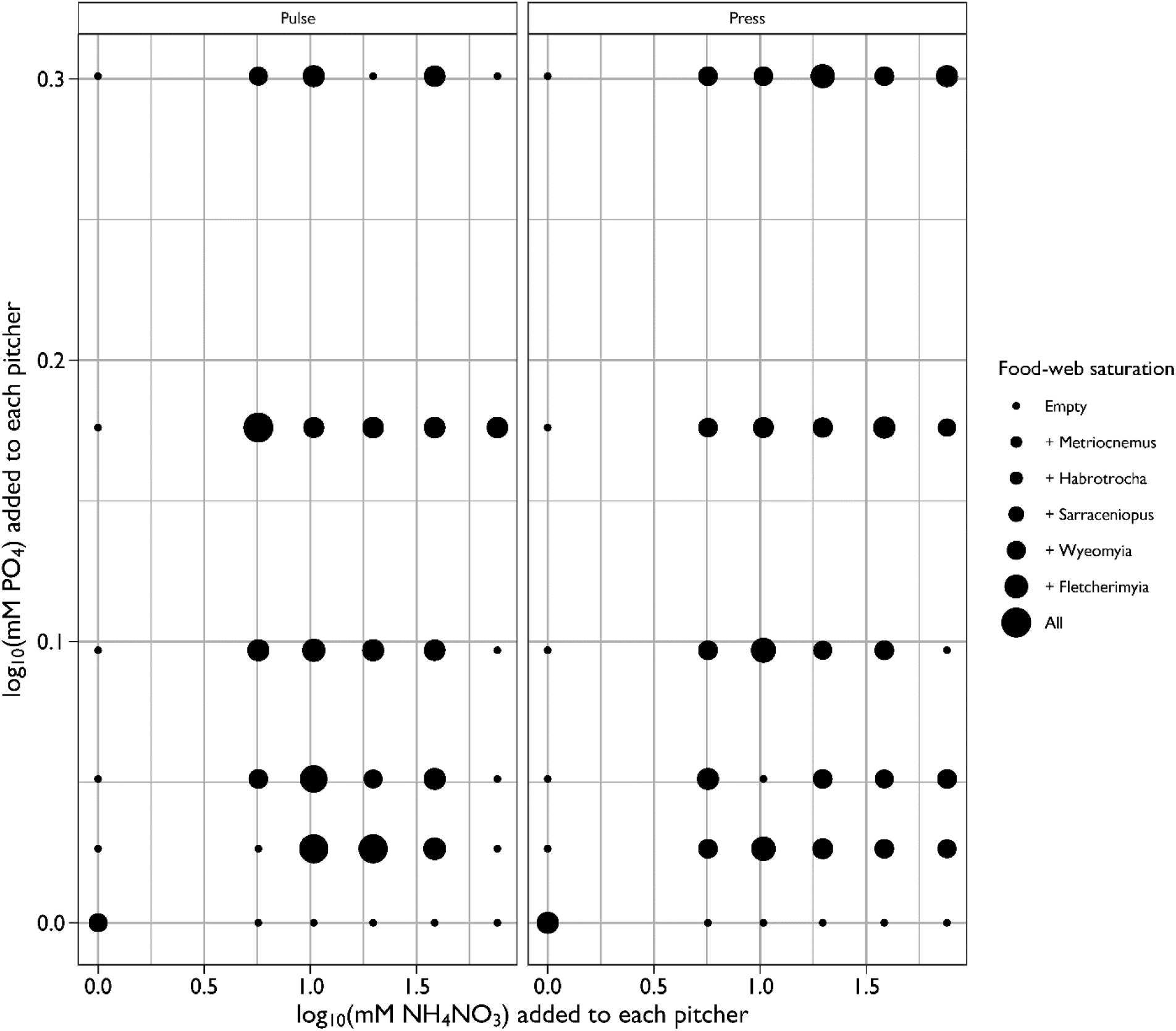
Response surface of food-web saturation (defined in Table 1) in pitchers fed one of 25 combinations of nitrogen (N) and phosphorus (P) or distilled water once, three days after the pitcher opened (the “pulse” experiment; left panel); or three days after the pitcher opened and every two weeks thereafter (the “press” experiment; right panel). This figure illustrates data from the first leaf of the season (fed on or beginning the week of 4 June 2001) at the mid-summer harvest (6 August 2001 for the pulse experiment, 23 July 2001 for the press experiment), when saturation was highest

Food-web saturation in the coincident press experiment also peaked at intermediate levels of N added (F_1,292_ = 3.7, P = 0.05) and was highest in mid-summer (F_1,292_ = 26.9, P < 0.01) in those pitchers that were produced earliest in the season (F_1,203_ = 4.1, P = 0.04) (Fig. 2). As in the pulse experiment, the amount of P added did not alter food web saturation (F_1,292_ = 0.4, P = 0.54), but in contrast to the results of the pulse experiment, there was no significant N × P interaction in the press experiment (F_1,292_ = 0.3, P = 0.58). We did not observe any pitcher mortality from the higher levels of N additions.

### Experiment 2

Inquilines colonized all the plants, but *Wyeomyia* was observed somewhat less frequently in pitchers with intact tubes than in pitchers with perforated tubes or no tubes at all (χ^2^_[2]_ = 28.5, P < 0.01; Table 2). Similarly, *Fletcherimyia* colonized only two pitchers with intact tubes and the added *Fletcherimyia* survived in only one pitcher with an intact tube (χ^2^_[1]_ = 20.6, P < 0.01; Table 2). Occurrences of the other macrobial inquilines were unrelated to tube treatment or *Fletcherimyia* addition (respectively χ^2^_[2]_ = 3.4, P = 0.18 and χ^2^_[1]_ = 0.03, P = 0.87; Table 2).

**Table 2.** Inquilines collected from 64 pitchers in the four-level full factorial experiment in which food webs were isolated from the plant with polyethylene tubes, inorganic and organic nutrients were added to assess bottom-up effects, and the top predator *Fletcherimyia fletcheri* was added to assess top-down effects. This table illustrates the effects on inquiline occurrence of the two factors—the polyethylene tubes and the addition of the top predator—that had significant effects on food-web saturation (Fig. 3). Values in each cell are the number of pitchers in which each inquiline was collected, pooled over inorganic and organic nutrient addition treatments and both harvests. The maximum possible number of occurrences for any given inquiline in a Tube × *Fletcherimyia* treatment combination is 32. Because cell entries within each column are occurrences summed over each Tube × *Fletcherimyia* treatment combination, all entries within a single row do not imply co-occurrences. Occurrences of the other macrobial inquilines within a tube treatment were unrelated to the addition of *Fletcherimyia*

|  |  | Number of pitchers in which inquiline occurred |  |  |  |  |
| --- | --- | --- | --- | --- | --- | --- |
| Tube | *Fletcherimyia* |  |  |  |  |  |
| treatment | treatment | *Metriocnemus* | *Habrotracha* | *Sarraceniopus* | *Wyeomyia* | *Fletcherimyia* |
| Intact | Added | 27 | 6 | 15 | 13 | 1 |
|  | Not added | 26 | 8 | 11 | 20 | 2 |
| Perforated | Added | 31 | 7 | 8 | 28 | 13 |
|  | Not added | 29 | 6 | 12 | 28 | 10 |
| None | Added | 24 | 2 | 11 | 28 | 14 |
|  | Not added | 28 | 1 | 6 | 24 | 6 |

Connections with the plant increased food-web saturation: food webs in pitchers without tubes or with perforated tubes were significantly more saturated than food webs in pitchers with intact tubes (Fig. 3, right panel; F_2,173_ = 8.4, P < 0.01). Removal of *Fletcherimyia* from plants led to an increase in food web saturation (Fig. 3, left panel; F_1,173_ = 13.2, P < 0.01), as this predator consumes both rotifers and *Wyeomyia* larvae. Neither additions of prey nor inorganic nitrogen (NH_4_NO_3_-N) alone altered food-web saturation (F_1,173_ = 0.8, P = 0.37; F_1,173_ = 0.0, P = 0.88, respectively). However, there was a nutrient × *Fletcherimyia* interaction (F_1,182_ = 4.9, P = 0.03): in the presence of *Fletcherimyia*, increasing NH_4_NO_3_ led to decreased saturation of the food web. In contrast, in the absence of *Fletcherimyia*, increasing NH_4_NO_3_ led to increased saturation of the food web. None of the other two-way, three-way, or four-way interaction terms was significant (P > 0.18, all cases).

**Fig. 3.**
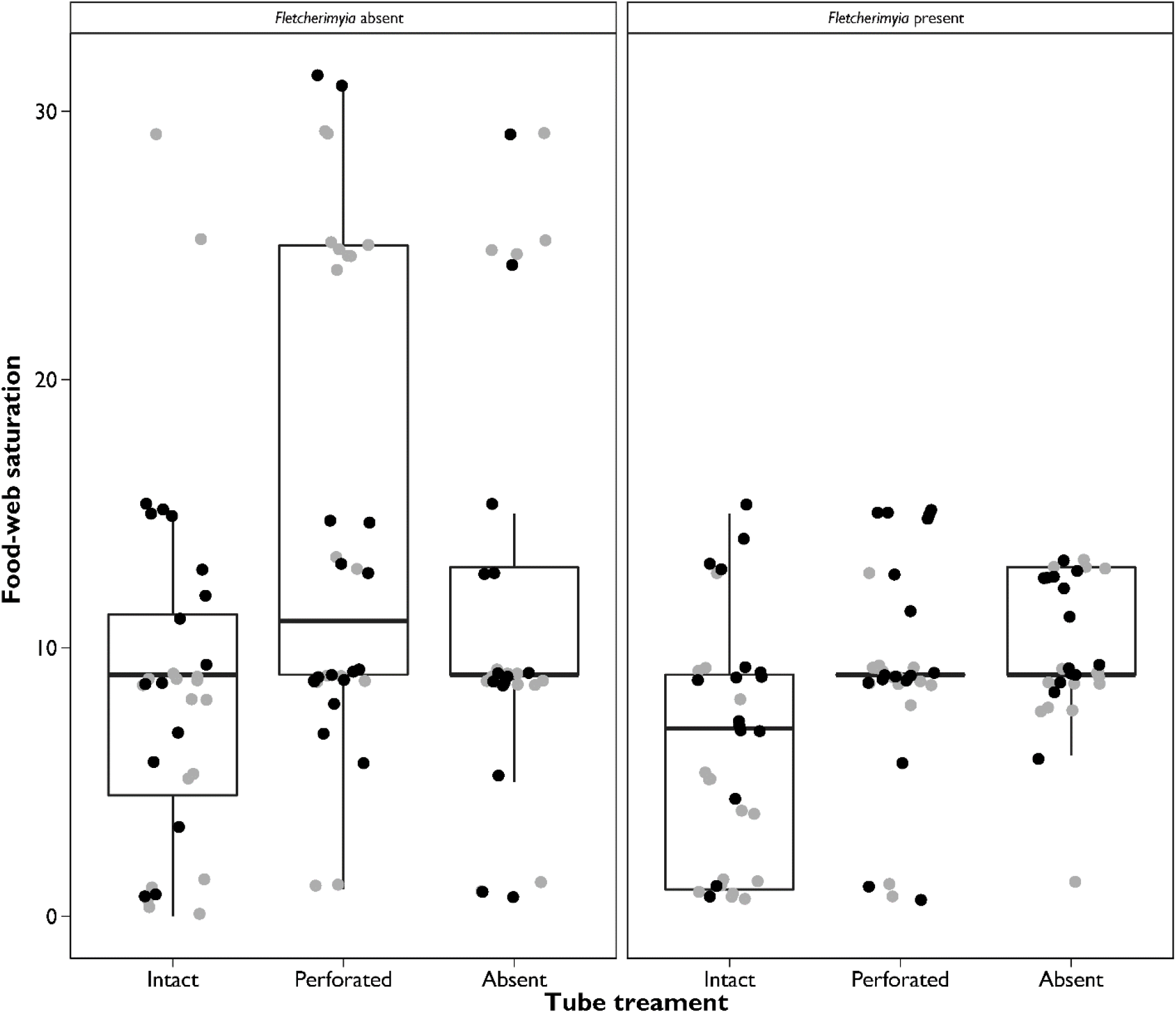
Effects of tube treatment manipulation of the top predator on food-web saturation in the *Sarracenia purpurea* macrobial food webs in the four-level tube × inorganic nitrogen × fruit-fly × *Fletcherimyia* manipulation. Data are pooled among nutrient and addition treatments, for which the main effects were not statistically significant (see main text for ANCOVA results). In each panel, the effects of the three levels of the inserted tubes (intact, perforated, absent) are ordered along the *x*-axis; the two different panels illustrate effects of the tube treatment without (left) or with (right) *Fletcherimyia* present in the pitcher. Box-and-whisker plots illustrate median (dark horizontal line), upper and lower quartiles (top and bottom of the box), and upper and lower hinges (length of vertical “whiskers” = 1.5×interquartile range). Individual points represent data from each pitcher harvested in mid-summer (grey points) or at the end of the growing season (black points).

## Discussion

Plant productivity, expressed as new biomass eaten by herbivores or shed litter decomposed by detritivores, exerts bottom-up control of most food webs (McCann 2012; Wokovich *et al.* 2014; Zou *et al.*, 2016). As phytotelmata (Maguire, 1971), plants produce habitat for aquatic food webs (Kitching, 2000, 2001; Ellison *et al.*, 2003; Srivastava *et al.*, 2004), but other effects of phytotelmata on food webs have been studied only rarely (Bradshaw and Creelman, 1984; Joel and Gepstein, 1985; Clarke and Kitching, 1995; Thornham *et al.*, 2011; Bazile *et al.*, 2012). The experiments described here illustrate that the phytotelmata of northern pitcher plants are not simply inert containers in which complex aquatic food webs assemble and thrive. Rather, the plant itself interacts with, and determines the strength of, the bottom-up and top-down controls of the food web.

Unlike pitchers in other species of *Sarracenia* that are covered by a reflexed ‘hood’, pitchers of *S. purpurea* are open to the sky and accumulate water from precipitation. This has two consequences. First, as pitchers accumulate water, they create the habitat for the inquiline food web. Second, any nutrients that accompany precipitation can enter the food web through bacterial mineralization and immobilization.

Independent of addition of inorganic nutrients, supplemental prey, or presence of top predators, the plant itself had a net positive effect on inquiline food-web saturation. Food webs in containers isolated from the plant were 10–20% less saturated than food webs in perforated tubes or unmanipulated controls of comparable size. This result agrees with previous observations of, and experiments with, *S. purpurea* and its inquilines, in which the plant oxygenated the pitcher fluid, maintaining homeostatic conditions for the inquiline food web and preventing a switch to anaerobic conditions (Bradshaw and Creelman, 1984; Joel and Gepstein, 1985; Sirota *et al.*, 2013).

Further addition of inorganic N, as is occurring through atmospheric deposition, increased food-web saturation at intermediate levels, but saturation declined as N addition was increased further (Fig. 2). We hypothesize that this result may reflect bottom-up effects of N on microbial productivity (Moorhead and Sinsabaugh, 2006), shifts in how *S. purpurea* uses different forms of N (Karagatzides *et al.*, 2009), or interactions between the plant and the bacteria that mineralize nitrogen for it (Butler *et al.*, 2008; Mouquet *et al.*, 2008). These studies, and others with excess prey addition show shifts in the macrobial food web or bacterial functional groups (see Northrop *et al.* 2021 for the latter) but no evidence of nutrient toxicity (*sensu* Camargo *et al.* 2005). Furthermore, N addition alters pitcher morphology, reducing pitcher size (volume) while increasing the area of its photosynthetic keel (Ellison and Gotelli, 2002; Bott *et al.*, 2008). Although smaller pitchers have less complex food webs (Gotelli and Ellison, 2006), this effect was not observed or measured here because we worked only with pitchers that were fully opened, not those that were produced after nutrients were added to early-season pitchers and that would have expressed morphological differences following nutrient translocation and use in new pitcher production (Butler and Ellison, 2007).

Top-down effects of predation in the inquiline food web of *S. purpurea* have been described repeatedly (*e.g.*, Addicott, 1974; Kneitel and Miller, 2002; Hoekman, 2007, 2011; Peterson *et al.*, 2008), but in our experiments, these effects were manifest only when the food web was isolated from the plant. Food webs in tubes rarely had predators (*Wyeomyia* or *Fletcherimyia*), as these Diptera would not oviposit in intact tubes and food webs without the predator *Fletcherimyia* were more saturated (up to the level below *Fletcherimyia*) than food webs with this predator. The interactions that we observed between top-down effects of *Fletcherimyia* and bottom-up effects of inorganic N additions are mirrored by modelled trade-offs in overall system nitrogen mineralization modulated by interactions between plants and mineralizing bacteria and between plants and higher trophic levels (Mouquet *et al.*, 2008; Ellison and Gotelli, 2021). Finally, the potential importance of digestive enzymes in mediating any of these produced by the plant remains unknown. When digestive enzymes are produced during pitcher development and at what concentrations remain open questions (*e.g.*, Gallie and Chang, 1977; Luciano and Newell, 2017). However, our results do illustrate that the pitcher itself does more than simply maintain a habitat for symbionts (*cf.* Luciano and Newell, 2017).

Overall, the results presented here together with those from other studies on pitcher-plant food webs illustrate that top-down and bottom-up effects on food-web structure are dependent on the habitat context in which they occur. Anthropogenic processes such as fossil-fuel combustion leading to atmospheric deposition of inorganic nutrients can alter habitat structure and shift control of food web structure from strictly bottom-up control to a mixture of top-down and bottom-up controls. These consequences are especially evident in habitats that themselves are living structures that respond to the same forcing processes at similar time scales. Plants such as *S. purpurea* that create phytotelmata are more than inert containers and their inhabitants are more than commensal inquilines. Rather, both the plant and the ‘inquilines’ are partners in complex networks of interactions. Our work and that of others illustrate clearly that the microecosystem created by *S. purpurea* can provide a model experimental system for exploring such interactions (Ellison and Gotelli, 2021).

## Acknowledgement

We thank Clarisse Hart, Callan Ordoyne, and Ali Rosenberg for significant assistance with the fieldwork, and two reviewers for excellent comments on an early version of the ms. This research was supported by NSF grants DEB 98-05722, 98-08504, 02-35128, and 02-34710 to AME, and permitted by the Massachusetts Department of Hatcheries.

## Notes

https://dx.doi.org/10.6073/pasta/3bd2c920cf431442d34227f3f5fc7c2a

### Competing Interest Statement

The authors have declared no competing interest.

### Summary of Updates

Revised version in response to reviews received. Resubmitted to American Midland Naturalist on 8 March 2021

